# Demonstration of short-term plasticity in the dorsolateral prefrontal cortex with theta burst stimulation: A TMS-EEG study

**DOI:** 10.1101/101097

**Authors:** Sung Wook Chung, Benjamin P. Lewis, Nigel C. Rogasch, Takashi Saeki, Richard H. Thomson, Kate E. Hoy, Neil W. Bailey, Paul B. Fitzgerald

## Abstract

**Objectives:** To examine the effects of intermittent TBS (iTBS) and continuous TBS (cTBS) on cortical reactivity in the dorsolateral prefrontal cortex.

**Methods:** 10 healthy participants were stimulated with either iTBS, cTBS or sham at F3 electrode. Single- and paired-pulse TMS and concurrent electroencephalography (EEG) were used to assess change in cortical reactivity and long-interval intracortical inhibition (LICI) via TMS-evoked potentials (TEPs) and TMS-evoked oscillations.

**Results:** Significant increases in N120 amplitudes (p < 0.01) were observed following iTBS over prefrontal cortex. Changes in TMS-evoked theta oscillations and LICI of theta oscillations were also observed following iTBS (increase) and cTBS (decrease). Change in LICI of theta oscillations correlated with change in N120 amplitude following TBS (r = −0.670, p = 0.001).

**Conclusions:** This study provides preliminary evidence that TBS produces direct changes in cortical reactivity in the prefrontal cortex. Combining TBS with TMS-EEG may be a useful approach to optimise stimulation paradigms prior to the conduct of clinical trials.

**Significance:** TBS is able to modulate cortical reactivity and cortical inhibition in the prefrontal cortex.

**Highlights:**

- Effects of iTBS and cTBS were studied in the DLPFC using TMS-EEG
- iTBS increased N120 amplitude, theta power and LICI of theta
- cTBS decreased theta power alone

## 1. Introduction

Repetitive transcranial magnetic stimulation (rTMS) is a non-invasive brain stimulation method capable of modulating the excitability of cortical circuits for extended periods of time and has been utilized in both research and clinical applications (Machado et al., 2013). A variety of rTMS methods have been shown to modulate cortical functioning, including high (>5 Hz) and low (~1 Hz) frequency forms of rTMS, which have demonstrated lasting, but divergent effects on cortical excitability (Grossheinrich et al., 2009, Cardenas-Morales et al., 2010, Guse et al., 2010, Esslinger et al., 2014). A more recently developed method of delivering rTMS involves the application of magnetic stimulation in specific frequency patterns, which are thought have greater efficacy in modulating cortical activity. Theta burst stimulation (TBS) involves the application of a burst of three pulses at 50 Hz repeated at 5 Hz. Continuous TBS (cTBS) employs a continuous series of bursts for a total of 40 seconds (600 pulses), and has been shown to decrease cortical excitability in the motor cortex measured using motor evoked potentials (MEPs). On the other hand, intermittent TBS (iTBS) involves the application of a 2-second train of TBS repeated every 10 seconds for a total of 192 seconds, and has been shown to increase MEP amplitude (Huang et al., 2005). TBS was originally developed based on the observations of natural firing patterns within the hippocampus, and the application of higher-frequency bursts (gamma) nested within lower-frequency rhythms (theta) resulted in reliable and robust long-term potentiation (LTP) (Larson et al., 1986). Rhythmic synchronization, in particular, coupling of theta and gamma frequencies has been shown to play a key role in the communication between neuronal networks, as well as in synaptic plasticity to promote learning and memory (Fell et al., 2011, Machado et al., 2013, Colgin, 2015, Lega et al., 2016). As such, this phase-locked firing pattern of stimulation appears to have effects on cortical excitability that are the equivalent, if not greater than those produced with traditional rTMS methods, and there is some evidence of longer-lasting aftereffect duration (Nyffeler et al., 2006, Yang et al., 2015). The vast majority of TBS studies have demonstrated reliable corticospinal excitability change following motor cortex stimulation (Wischnewski et al., 2015). However, recent studies with larger sample sizes have shown no group-level effect of TBS (Hamada et al., 2013, Hinder et al., 2014), and a meta-analysis has revealed publication bias (Chung et al., 2016), suggesting a degree of inter-individual variability. Furthermore, little is known about the effects of TBS on cortical excitability/inhibition outside the motor cortex. This is important as there is increasing interest in the use of TBS as a therapeutic tool for disorders such as depression (Li et al., 2014, Plewnia et al., 2014, Bakker et al., 2015, Chistyakov et al., 2015, Chung et al., 2015a, Cheng et al., 2016) and schizophrenia (Demirtas-Tatlidede et al., 2010, Brunelin et al., 2011, Hasan et al., 2015), where dorsolateral prefrontal cortex (DLPFC) is the primary target for treatment. In developing clinical applications using TBS, there is a need to explore the cortical excitability effects of TBS in non-motor cortical regions. Establishing that changes in cortical excitability due to TBS can be detected with TMS-EEG will also allow the use of these approaches in preclinical studies investigating the optimal methods to induce changes in cortical activity.

Measuring TMS-evoked activity with EEG can be used to assess changes in cortical properties before and after an intervention (for review, see (Chung et al., 2015b)), and allows investigation beyond motor cortex. A number of studies have previously described TMS-evoked potentials (TEPs) found at latencies of 40, 60, 100 and 200 ms post a single stimulus and attempted to understand how these TEPs relate to aspects of cortical function (Rogasch et al., 2013a, Bortoletto et al., 2015, Gosseries et al., 2015). For example, TEPs at latencies of ~100 ms (N120) has been associated with cortical inhibitory processes (Bikmullina et al., 2009, Rogasch et al., 2013a, Premoli et al., 2014a). However, the origin of other TEP components is still largely unknown.

The aim of this study was to examine the effects of iTBS and cTBS on cortical reactivity in the dorsolateral prefrontal cortex (DLPFC), a brain region relevant to the treatment of a number of neuropsychiatric disorders including depression, schizophrenia and movement disorders (Spronk et al., 2008, Cardenas-Morales et al., 2010, Leonard et al., 2013). Healthy subjects were stimulated in three separate sessions with either iTBS, cTBS or sham stimulation with the EEG responses to single and paired TMS pulses used to assess cortical reactivity and inhibition respectively, pre-and post-TBS. We hypothesized that an iTBS and cTBS protocol would increase and decrease cortical reactivity respectively with respect to sham condition. In addition, we hypothesised that TBS-induced changes would be observed in the theta and gamma band oscillations, as these frequencies are targeted by TBS.

## 2. Materials and Methods

### 2.1 Participants

10 right-handed participants (mean age 31.3 ± 9.3 years, range 21-51, 4 female) completed all 3 testing sessions. A number of state-dependent factors including the effect of medications, and substances such as alcohol and caffeine, can affect a participant’s response to TBS (Silvanto et al., 2008). Participants were asked to refrain from the consumption of alcohol and caffeine prior to the experiment and participants taking psychotropic medications were excluded from the study.

Four participants had partaken in previous TMS experiments but none in the month prior to participation. All participants gave informed consent prior to the experiment and were screened with the Mini International Neuropsychiatric Interview (MINI) in order to exclude psychopathology (Sheehan et al., 1998). The Alfred Hospital and Monash University Human Research and ethics committee approved the experiment.

### 2.2 Procedure

This study consisted of the recording of EEG responses to 50 single and paired TMS pulses both before and after one type of TBS at each session. Each participant attended for three sessions with at least one week apart, and the sessions were pseudorandomized and counterbalanced in administering either cTBS, iTBS or sham stimulation. Participants were required to keep their eyes open throughout the experiment and interaction with participants was limited during testing.

#### 2.2.1 TMS

TMS was targeted to the left DLPFC throughout the experiment utilising the 10/20 method of electrode placement. It has been shown that DLPFC is approximately at the midpoint between the F3 and AF3 electrode position in 10/20 system, but slightly closer to F3 (Fitzgerald et al., 2009), and therefore the centre of the coil was placed at F3 electrode. In addition, the coil was positioned at 45 degrees to the midline as it has shown to produce the strongest stimulation in the DLPFC (Thomson et al., 2013). A Magstim 200 stimulator with a figure of eight coil (Magstim Ltd, Withland, Wales, UK) was used for single and paired pulse stimulation pre- and post-TBS using monophasic pulses (posterior-anterior current direction in the underlying cortex). TBS was delivered with a single MagVenture B-65 fluid-cooled coil (MagVenture, Copenhagen, Denmark) using biphasic pulses (antero-posterior to postero-anterior current direction in the underlying cortex). Stimulus intensity was set relative to resting motor threshold (rMT) obtained from the left motor cortex, which was identified via Ag/AgCl EMG electrodes attached to the first dorsal interosseous (FDI) muscle. The rMT was used to calibrate and normalise TMS coil output energy for inter- and intra-individual physiological variability, and was measured at the beginning of each session separately for each TMS machine. The EEG cap was mounted, and the rMT was determined as the minimum intensity required to evoke at least 3 out of 6 MEPs > 0.05 mV in amplitude (Conforto et al., 2004).

Participants received 50 single (5s interval ± 10% jitter) and 50 paired pulses (100 ms interval) over the DLPFC at 120% rMT before and after TBS. cTBS, iTBS or sham (depending on the session) was delivered at 80% rMT: cTBS-a burst of 3 pulses of stimulation given at 50Hz repeated every 5Hz for a total of 40s, iTBS-a 2s train of TBS repeated every 10s for a total of 192s. Sham stimulation was applied using the iTBS protocol, with the coil orientated at 90 degrees to the scalp so the magnetic field would be delivered tangentially away from the scalp. Each condition delivered a total of 600 pulses.

#### 2.2.2 EEG recording

EEG responses to TMS pulses were recorded with a 64-channel TMS-compatible EEG cap and a TMS-compatible EEG amplifier (SynAmps2, EDIT Compumedics Neuroscan, Texas, USA). Recordings were obtained from 39 electrodes positioned around the scalp (AF3, AF4, F7, F5, F3, F1, Fz, F2, F4, F6, F8, FC5, FC3, FC1, FCz, FC2, FC4, FC6, T7, C5, C3, C1, Cz, C2, C4, C6, T8, P7, P5, P3, P1, Pz, P2, P4, P6, P8, O1, Oz, O2). To monitor eye blinks, electrooculography (EOG) recordings were obtained from 4 Ag/AgCl electrodes, two positioned above and below the left eye and two positioned lateral to the outer canthus of both eyes.

Electrodes were online referenced to CPz and grounded to POz. Signals were amplified (1,000 x), low-pass filtered (2,000 Hz) and recorded on a computer for offline analysis. The high acquisition rate (10,000 Hz), large operating range (± 200 mV) and DC coupling of the EEG amplifier allows recording of the TMS artefact without amplifier saturation. Electrode impedance levels were kept at <5 kΩ throughout the experiment.

The loud auditory click produced by the TMS coil often produces an auditory EEG response. This was in part controlled by the application of white noise through earphones during TMS/TBS of the experiment.

### 2.3 EEG data preprocessing

TMS-EEG data were analysed offline using EEGLAB (Delorme et al., 2004), FieldTrip (Oostenveld et al., 2011) and custom scripts on the Matlab platform (R2015b, The MathWorks, USA). Data were epoched around the test TMS pulse (−1,000 to 1,000 ms), baseline corrected with respect to the TMS-free data (−500 to −110 ms), and data between - 5 and 10 ms (around the large signal from the TMS pulse) were removed and linearly interpolated. This baseline period was chosen so that the baseline period is equivalent between single and paired conditions. Both pre- and post-TMS epoched data for each condition were concatenated and analysed concurrently so that rejected components were removed in both sets of data to avoid bias. Data were downsampled to 1,000 Hz and visually inspected to remove epochs containing muscle artefact or excessive noise. An average of 5.4 trials was rejected in the cTBS condition, 6.2 trials in the iTBS condition and 2.8 trials in the sham condition across both pre and post conditions. Bad channels (e.g. disconnected) were then removed. An initial round of independent component analysis (ICA) using the symmetric FastICA algorithm with nonlinearity ‘tanh’ was applied to remove the remainder of the muscle artefact (Korhonen et al., 2011). The remaining tail of TMS-evoked muscle artefact components was identified by the size, topography and properties of the waveform, which have been shown to be consistent with scalp muscle activation (Rogasch et al., 2013b), and was subtracted from the data. All data were bandpass filtered (second-order, zero-phase, Butterworth filter, 1-100Hz & notch at 50 Hz) and epochs were inspected again to remove any anomalous activity in the signal such as excessive muscle activity from jaw-clenching. The data were then submitted to the FastICA algorithm again. Identification and removal of additional artefactual components were based on a previous study (Rogasch et al., 2014), where blink artefacts, decay artefacts and other noise-related artefacts were removed. Removal of auditory evoked potentials described in the aforementioned study was not adopted in the current study as partial noise-masking was applied. Removed channels were interpolated, and data were re-referenced to common average reference. Finally, remaining epochs from each trial within a block were averaged.

#### 2.3.1 TMS-evoked potentials (TEPs)

TEPs were analysed using two different methods. A region of interest (ROI) analysis of the average of 4 electrodes (F3, F1, FC3, FC1) was conducted to access the local effects of TBS. These electrodes were close to the point of stimulation and sat above left prefrontal cortex. The average of these electrodes also adequately represents all of the major peaks (Fig. 1). To ensure the spatial distribution was properly captured, cluster-based statics were used to evaluate global effect of TBS across the scalp. TEPs were compared before and after each type of TBS and compared across different type of TBS. Four peaks (N40, P60, N120 and P200) were chosen based on previous TMS-EEG studies on the DLPFC (Rogasch et al., 2014, Rogasch et al., 2015). TEP peaks were selected within pre-defined time windows for N40 (30 – 50 ms), P60 (55 – 75 ms), N120 (90 – 140 ms) and P200 (160 – 240 ms) at the ROI electrodes. The peak amplitude was then calculated as the average signal between ± 5 ms of the selected peak latency. The same latency windows were used for global scalp analysis.

**Figure 1.**
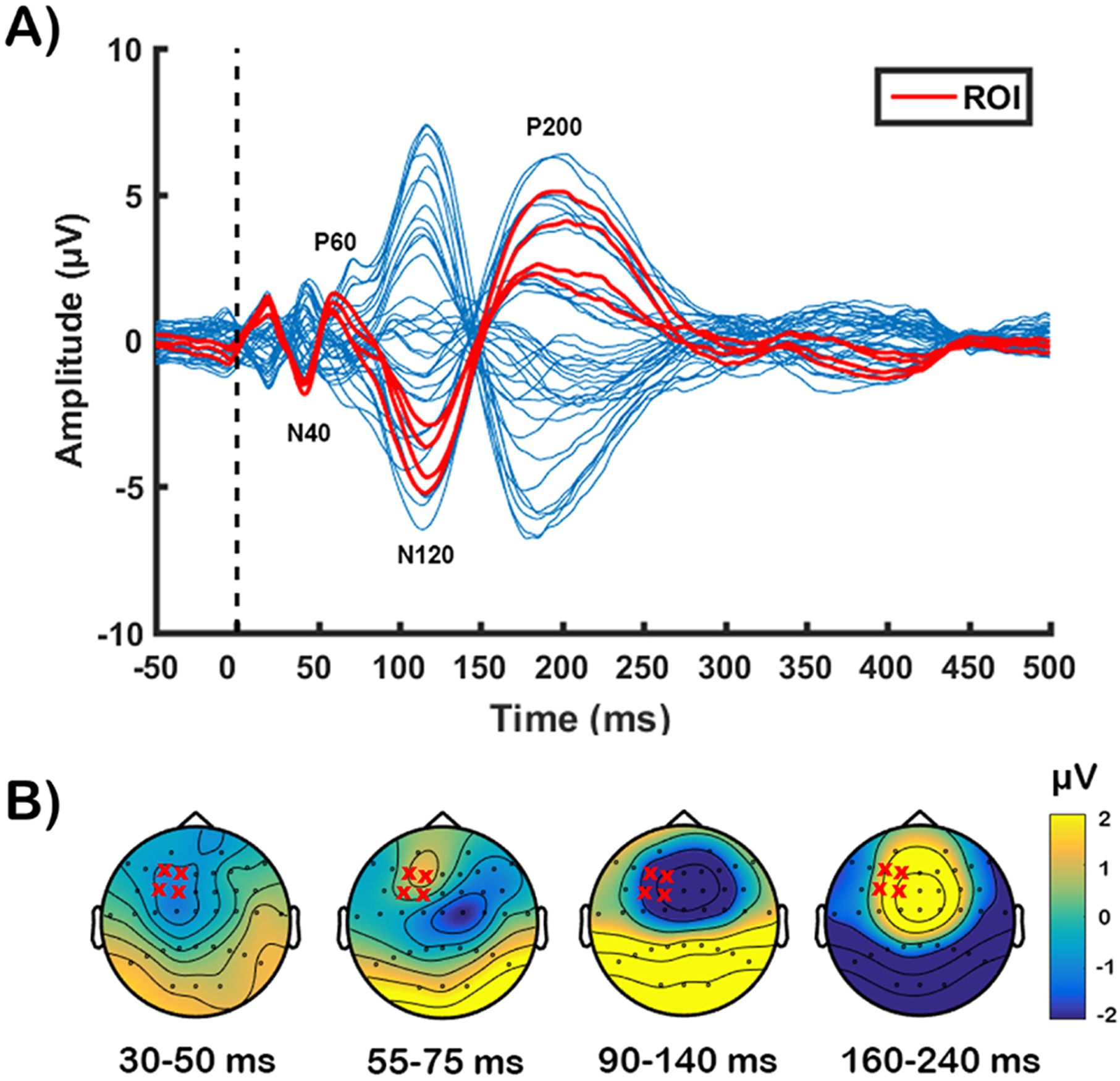
TMS-evoked potential following single-pulse stimulation over left dorsolateral prefrontal cortex before theta-burst stimulation (averaged across participants). (A) Butterfly plot from all electrodes with major peaks (N40, P60, N120, P200) indicated in text. The red lines indicates the waveform obtained from 4 electrodes (F3, FC3, F1, FC1) around the stimulation site. (B) Voltage distribution for each peak of interest averaged across time indicated below. Red ‘x’ mark indicates the 4 electrodes chosen.

For the paired-pulse condition, the single-pulse TMS data was subtracted from the conditioning pulse to account for ongoing effects of the conditioning pulse on the test pulse TEP (Rogasch et al., 2015). For LICI of TEPs, difference between unconditioned (single test pulse) and conditioned (paired test pulse) TEPs were normalised to the value of the overall size of the TEPs (30 – 240 ms) using the following formula for each peak of interest:

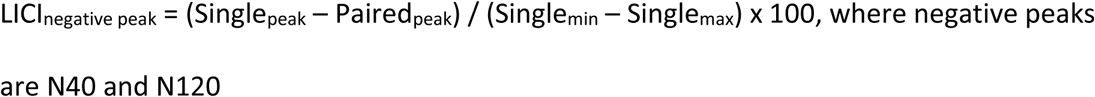

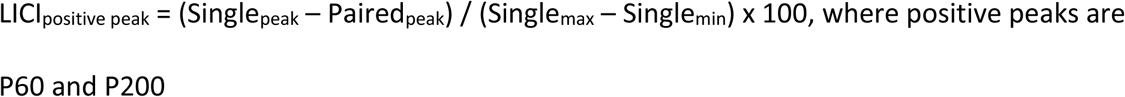

Normalisation of the data within a uniform range is necessary to consistently compare differences between single- and paired-pulse responses from peaks of different amplitudes.

#### 2.3.2 TMS-evoked oscillations

TMS-evoked oscillatory power was calculated by converting TEPs into the frequency domain using Morlet wavelet decomposition (3.5 oscillation cycles with steps of 1 Hz between 5 Hz and 70 Hz) on each trial for each electrode. Oscillatory power was then averaged over all trials, maintaining both evoked and induced oscillations. Normalised power was obtained through division of power by a mean baseline value (−650 to −350 ms). ROI analysis of the average of 4 electrodes (F3, F1, FC3, FC1) and global scalp analysis was again used to evaluate the effect of TBS on TMS-evoked oscillations. Oscillations were averaged across frequency bands of interest (theta [5 – 7 Hz] and gamma [30 – 70 Hz]) and across time (50 – 250 ms for theta, 25 – 125 ms for gamma). Theta and gamma frequencies were chosen for the direct reflection of the stimulation parameters, and the interaction of the two frequencies are implicated in synaptic plasticity (Nyhus et al., 2010, Zheng et al., 2015). For LICI of TMS-evoked oscillations, the difference between unconditioned and conditioned oscillations in theta, and gamma bands were normalised to the mean power across all frequency band (5 – 70 Hz) using the following formula:

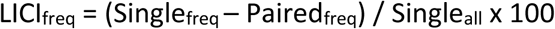

### 2.4 Statistical analysis

Statistical analysis was performed in SPSS (Version 22) and Matlab. For the ROI analysis, when data did not meet the requirement for normality (Shapiro-Wilk test), data were winsorised by setting extreme values to the corresponding adjacent 5^th^ or 95^th^ percentile value (Wilcox, 1997). A total of 2.08% of the data were winsorised for single-pulse TEPs and 2.5% of data for LICI of TEPs. A total 1.67% data were winsorised for each TMS-evoked oscillations and LICI of TMS-evoked oscillations. Repeated measures 2 [time (pre and post)] x 3 [stimulation conditions (iTBS, cTBS and Sham)] ANOVAs were performed on ROI analysis of TEP amplitudes and TMS-evoked oscillatory power. Post hoc t – tests were conducted with a Bonferroni correction applied to further explore the significant main effects of stimulation.

For global scalp analysis, non-parametric cluster-based permutation statistics were used (Oostenveld et al., 2011). To assess whether each TBS condition altered cortical reactivity over time, differences between pre- and post-TBS measures were compared. To assess whether TBS conditions differentially altered cortical reactivity, relative changes in TEP amplitudes and TMS-evoked oscillatory power following TBS (post – pre) between conditions were compared. Monte Carlo *p*-values were calculated from 5,000 randomizations and clusters were defined by two or more neighbouring electrodes with t-statistics at a given time or frequency point exceeding a threshold of *p* < 0.05 (dependent t-test).

It is recommended to analyse time-frequency data with regard to all of its possible dimensions (van Ede et al., 2016), and therefore, initial cluster-based analyses were performed including all data dimensions (time x frequency x space) for global analysis of TMS-evoked oscillations. Given our strong a priori hypotheses regarding the modulation of theta and gamma oscillations by TBS, we also applied more targeted analyses on these frequency bands (Averaged across 5 – 7 Hz, 50 – 250 ms for theta power, 30 – 70 Hz, 25 – 125 ms for gamma power).

To assess the relationship between changes in single and paired pulse markers of inhibition, Pearson’s correlations were used to compare stimulation-induced change in N120 amplitude with the change in LICI of TEPs and LICI of TMS-evoked oscillations.

## 3. Results

### *3.1.* Single-Pulse TMS

Single-pulse TMS over DLPFC resulted in a series of negative and positive peaks before and after TBS including N40, P60, N120 and P200 (See Table 1). There was no significant difference in the latency among different TBS conditions (all *p* > 0.05), and in the number of subjects in which peaks were found (iTBS: 10, cTBS: 10, sham: 9.75 ± 0.5). When peaks were not found, data were extracted from the pre-defined latencies (eg. 40 ms for N40). Data were re-analysed excluding one subject for the analyses associated with N40, however, results did not change.

**Table 1.** Latencies of each peak before and after each stimulation condition

|  | iTBS [mean (SD)] |  | cTBS [mean (SD)] |  | Sham [mean (SD)] |  |
| --- | --- | --- | --- | --- | --- | --- |
|  | T0 | T1 | T0 | T1 | T0 | T1 |
| **N40** | 37.7 ($\pm 6.91$) | 40.5 ($\pm 9.24$) | 41.8 ($\pm 8.94$) | 42.9 ($\pm 10.80$) | 36.4 ($\pm 8.42$) | 39.0 ($\pm 7.41$) |
| **P60** | 64.8 ($\pm 8.11$) | 65.5 ($\pm 11.18$) | 63.1 ($\pm 8.46$) | 63.7 ($\pm 8.47$) | 58.1 ($\pm 8.12$) | 60.9 ($\pm 6.19$) |
| **N120** | 118.5 ($\pm 10.29$) | 117.3 ($\pm 9.23$) | 115.0 ($\pm 6.60$) | 116.2 ($\pm 6.37$) | 115.2 ($\pm 10.48$) | 113.3 ($\pm 8.59$) |
| **P200** | 208.6 ($\pm 19.58$) | 213.5 ($\pm 21.21$) | 197.5 ($\pm 24.52$) | 200.4 ($\pm 23.06$) | 204.5 ($\pm 25.27$) | 202.5 ($\pm 29.08$) |

#### 3.1.1 ROI analysis of the effect of TBS on TEPs

Grand average TEP waveforms are illustrated in Fig. 2. Two-way repeated measures ANOVA yielded a trend in the main effect of time (*F*_1,9_ = 4.244, *p* = 0.069) and a significant interaction (*F*_2,18_ = 6.862, *p* = 0.006) in N120 amplitude. No significant main effect of condition was found (F2,18 = 2.565, *p* = 0.105). In order to investigate the interaction effect, a series of one-way ANOVA was performed across conditions at each time point (pre-and post-TBS), as well as paired t-tests across time for each condition. No significant main effect was observed at pre-TBS (*F*_2,18_ = 1.124, *p* = 0.350), but a significant main effect was found at post-TBS (*F*_2,18_ = 5.079, *p* = 0.018). Post hoc pairwise comparison revealed that N120 amplitude was significantly larger following iTBS compared to sham (*p* = 0.015), however there was no significant differences between iTBS and cTBS (*p* = 0.157), nor between cTBS and sham (*p* = 1.000). Across time within conditions, paired t-tests showed significant increase in N120 amplitude following iTBS (*t*_(9)_ = −3.297, *p* = 0.009), but no significant change was observed following cTBS (*t*_(9)_ = 0.182, *p* = 0.860) or sham *t*_(9)_ = −0.777, *p* = 0.457). No other peaks showed any significant main effects or interaction (all *p* > 0.05).

**Figure 2.**
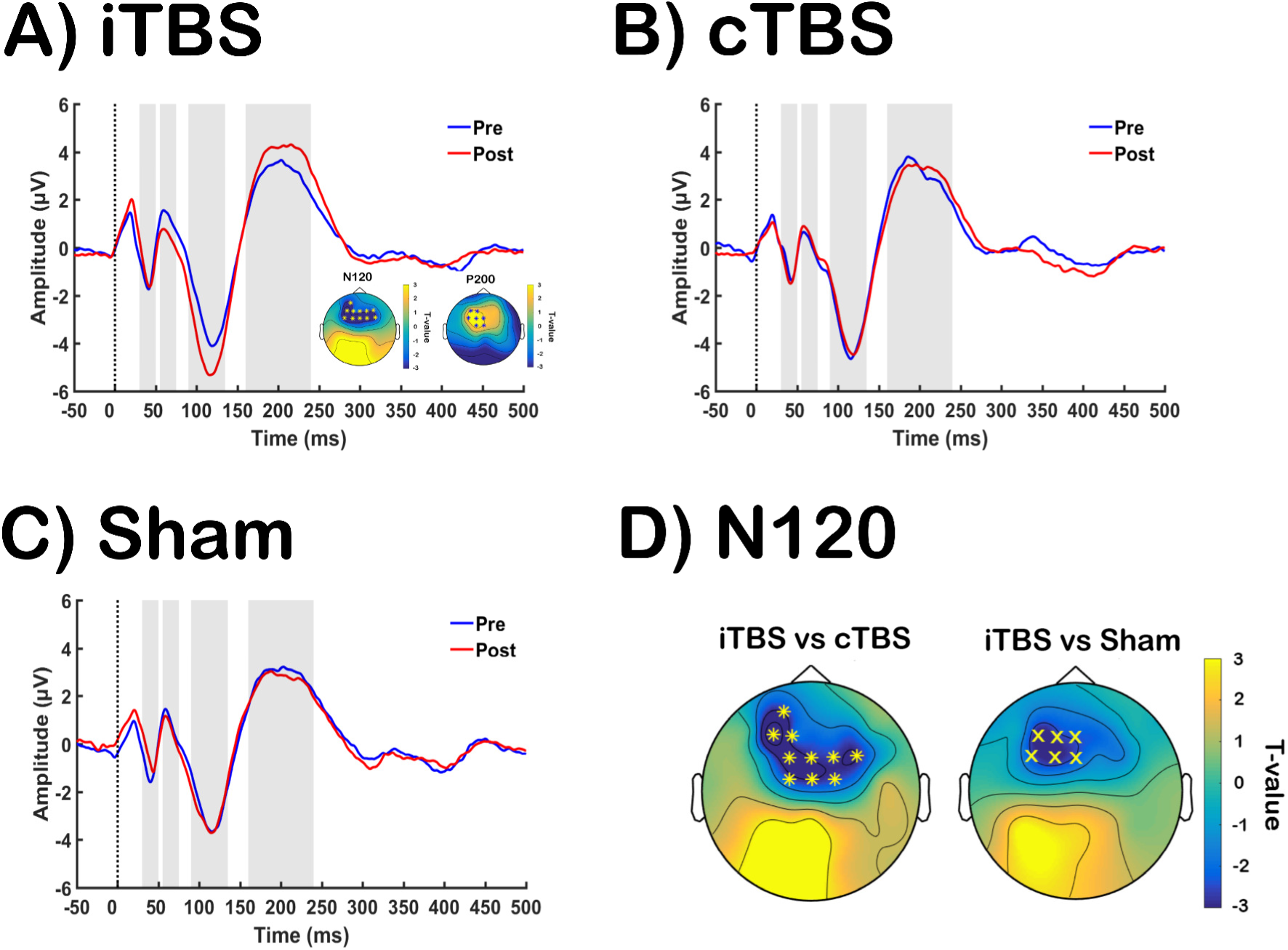
Assessment of transcranial magnetic stimulation (TMS)-evoked potentials (TEPs) before and after each stimulation condition [A: intermittent theta-burst stimulation (iTBS); B: continuous theta-burst stimulation (cTBS); C: Sham]. Grand average TEP waveforms pre- (blue) and post-TBS (red) for each stimulation conditions, with significant global differences illustrated in topoplots (post more negative (blue) or positive (yellow) than pre). (D) Comparison of different TBS induced N120 amplitude change at global level. Asterisks and ‘X’s on topoplots indicate significant clusters between comparisons (cluster-based statistics, \**p* < 0.01, ^X^*p* < 0.05).

#### 3.1.2 Global scalp analysis of the effect of TBS on TEPs

Over time, cluster-based statistics across space showed one significant negative cluster (p = 0.0008; post signal more negative than pre) at N120, indicating the N120 was more negative following iTBS, and one positive cluster (*p* = 0.0098; post signal more positive than pre) at P200, indicating P200 was more positive following iTBS. Topographical representation revealed a difference in N120 amplitude at the site of stimulation and contralaterally, while the difference in P200 amplitude was close to the stimulation site alone (Fig. 2A). There were no significant differences in other peaks across the scalp (all *p* > 0.05). No significant differences were observed between pre and post TEPs following cTBS (Fig. 2B) and sham (Fig. 2C) for any peaks (all *p* > 0.05).

In order to evaluate the differences between conditions, TBS-induced changes (post-pre) in TEP amplitude were calculated and compared. One significant negative cluster for the change in N120 (Δ N120) was found between iTBS and cTBS (*p* = 0.0002; iTBS more negative than cTBS), and between iTBS and sham (*p* = 0.017; iTBS more negative than sham). Topographical representation showed the differences were at the site of stimulation for both comparisons, but also contralaterally for the former (Fig. 2D). One positive cluster was observed with each comparison for the change in P200 (Δ P200) [iTBS vs cTBS (*p* = 0.009); iTBS vs sham (*p* = 0.005)], suggesting larger Δ P200 amplitude with iTBS compared to both cTBS and sham. The differences were observed at the stimulation site for cTBS, and frontocentral site for sham.

#### 3.1.3 ROI analysis of the effect of TBS on TMS-evoked oscillations

Two-way repeated measures ANOVA yielded no significant effect of time (*F*_1,9_ = 0.033, *p* = 0.859), but a trend in the main effect of condition (*F*_2,18_ = 3.211, *p* = 0.064) and a significant interaction (*F*_2,18_ = 7.857, *p* = 0.004) in TMS-evoked theta power. To assess the interaction effect, a series of one-way ANOVA was performed across conditions at each time point (pre- and post-TBS), and paired t-tests were conducted across time for each condition. No significant main effect was observed at pre-TBS (*F*_2,18_ = 2.529, *p* = 0.108), but a significant main effect was found at post-TBS (*F*_2,18_ = 6.876, *p* = 0.006). Post hoc pairwise comparison revealed that theta power was significantly larger following iTBS compared to sham (*p* = 0.046), and a trend was seen between iTBS and cTBS (p = 0.069). However, there was no significant difference between cTBS and sham (*p* = 1.000). Across time within conditions, paired t-tests showed a significant increase in theta power following iTBS (*t*_(9)_ = 2.416, *p* = 0.039), a significant decrease in theta power following cTBS (*t*_(9)_ = 2.377, *p* = 0.041), but no significant change was observed following sham stimulation (*t*_(9)_ = 0.007, *p* = 0.994) (Fig. 3A). No significant main effects or interactions were observed in TMS-evoked gamma power (p >0.05)

**Figure 3.**
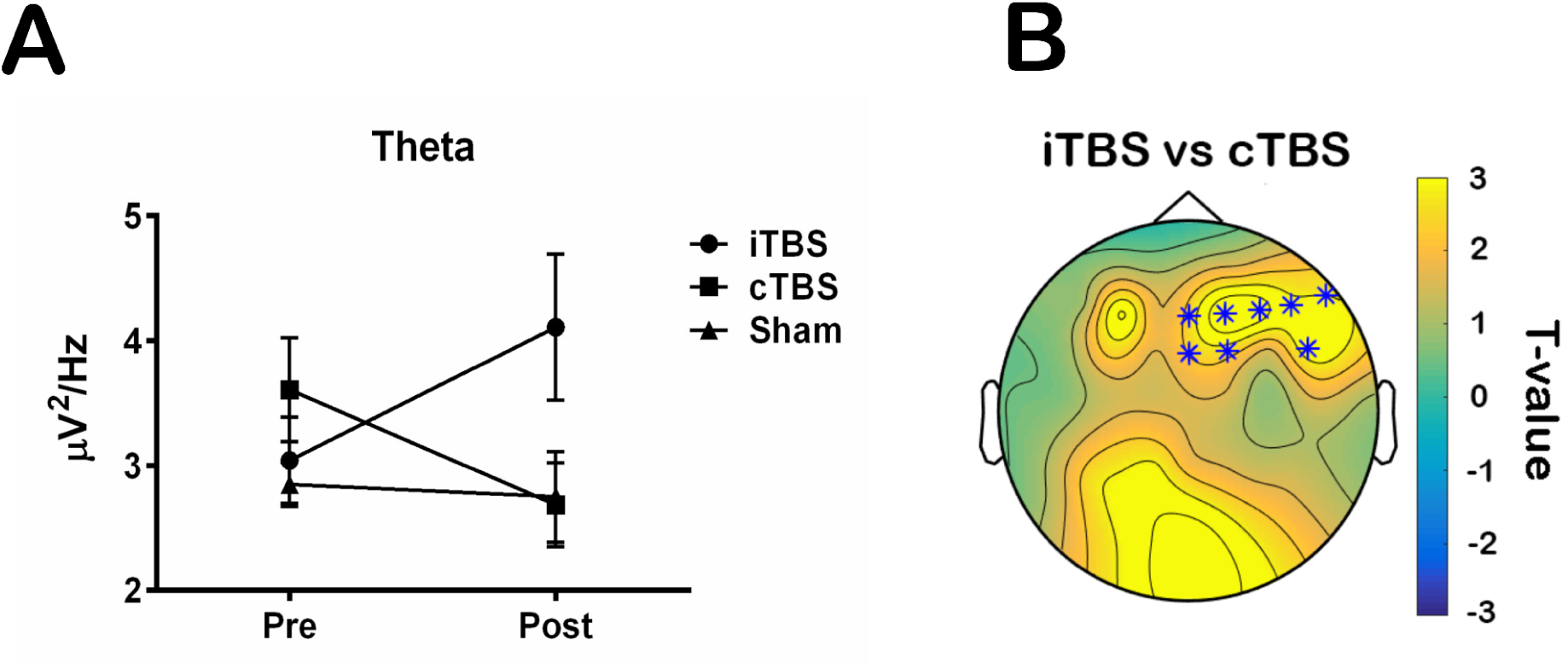
Comparison of transcranial magnetic stimulation (TMS)-evoked oscillations in TBS-induced changes [intermittent theta-burst stimulation (iTBS); continuous theta-burst stimulation (cTBS); Sham]. (A) TBS-induced theta power change over time among each stimulation condition at the region of interest and (B) comparison of TBS-induced theta power change across the scalp between iTBS and cTBS condition. Asterisks on topoplot indicate significant clusters between Δ theta post-iTBS and post-cTBS (cluster-based statistics, \**p* < 0.01)

#### 3.1.4 Global analysis of the effects of TBS conditions on TMS-evoked oscillations

To assess the effect of TBS across the scalp in the frequency domain, TMS-evoked oscillations were compared across space using cluster-based statistics. No significant clusters were found in any frequency band (all *p* > 0.05) within any stimulation condition across time. In order to evaluate the differences in TMS-evoked oscillations among stimulation conditions, TBS-induced changes (post – pre) were calculated and compared. One significant positive cluster in the change in theta power (Δ theta) was found between iTBS and cTBS (*p* = 0.008; Δ theta post-iTBS larger than post-cTBS; Fig. 3B) on the contralateral site of stimulation. No other significant cluster was found between other conditions.

### 3.2 Paired-pulse TMS (LICI)

#### 3.2.1 The effect of TBS on LICI of TEPs

The grand average TEP waveforms of single and paired-pulses are illustrated in Fig. 4. Two-way repeated measures ANOVA at ROI electrodes showed no significant main effects in LICI of TEPs in any peak (all *p* > 0.05). Globally, no significant cluster was found between pre- and post-TBS, and between conditions in any peak (all *p* > 0.05).

**Figure 4.**
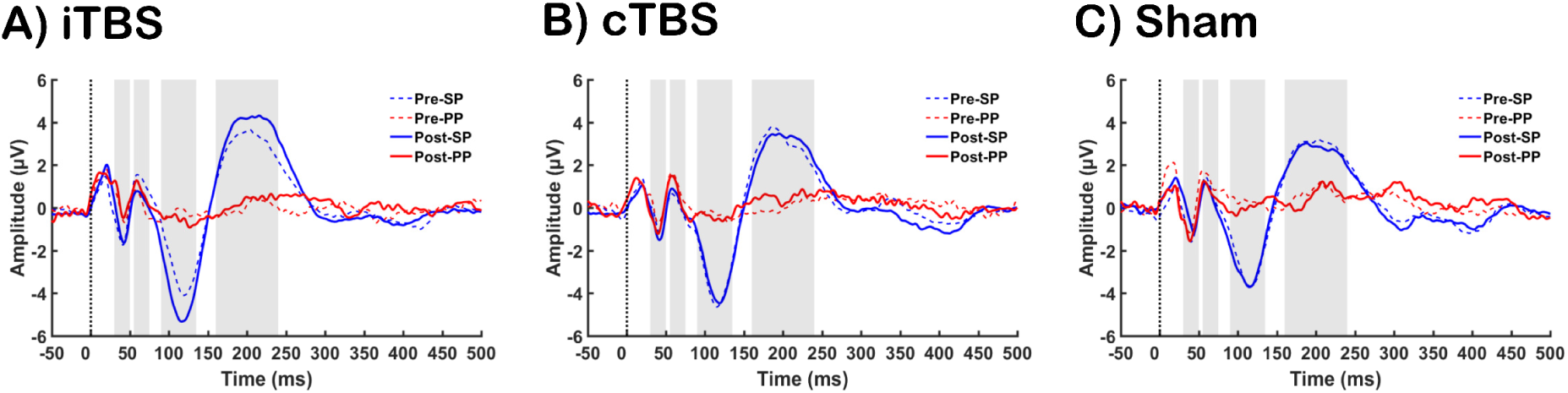
Assessment of long-interval intracortical inhibition (LICI) before and after each stimulation condition [A: intermittent theta-burst stimulation (iTBS); B: continuous thetaburst stimulation (cTBS); C: Sham], at F3 electrode. Grand average transcranial magnetic stimulation (TMS)-evoked potential (TEP) waveforms illustrate both single-pulse (SP; blue) and paired-pulse (PP; red) before (dotted-line) and after (solid-line) TBS.

#### 3.2.2 ROI analysis of the effect of TBS on LICI of TMS-evoked oscillations

Two-way repeated measures ANOVA yielded no significant effect of time (*F*_1,9_ = 0.010, *p* = 0.922), but a significant effect of condition (*F*_2,18_ = 5.421, *p* = 0.014) and a significant interaction (*F*_2,18_ = 8.156, *p* = 0.003) in LICI of theta power. To investigate the interaction effect, a series of one-way ANOVA was performed across conditions at each time point (pre-and post-TBS), and paired t-tests were conducted across time for each condition. No significant main effect was observed at pre-TBS (*F*_2,18_ = 2.740, *p* = 0.110), but a significant main effect was found at post-TBS (*F*_2,18_ = 6.644, *p* = 0.001). Post hoc pairwise comparison revealed that LICI of theta power was significantly larger following iTBS compared to sham (*p* = 0.007), and compared to cTBS (*p* = 0.029). However, no significant differences were observed between cTBS and sham (*p* = 1.000). Across time within conditions, paired t-tests showed a significant increase in LICI of theta power following iTBS (*t*_(9)_ = 3.030, *p* = 0.014), but no significant changes were observed following cTBS (*t*_(9)_ = −1.652, *p* = 0.133) or sham stimulation (*t*_(9)_ = 0.243, *p =* 0.814). No significant main effect was observed with LICI of gamma power (*p* > 0.05).

To assess whether Δ N120 following TBS was associated with the change in LICI (Δ LICI) of theta frequency band, correlation analysis was performed. Pearson’s correlation showed the change in N120 amplitude with TBS significantly correlated with change in LICI strength in the theta band (*r* = −0.670, *p* =0.001; Fig 5A), suggesting a larger increase in the negativity of the N120 amplitude corresponds to a larger increase in inhibition of theta frequency oscillations following TBS.

**Figure 5.**
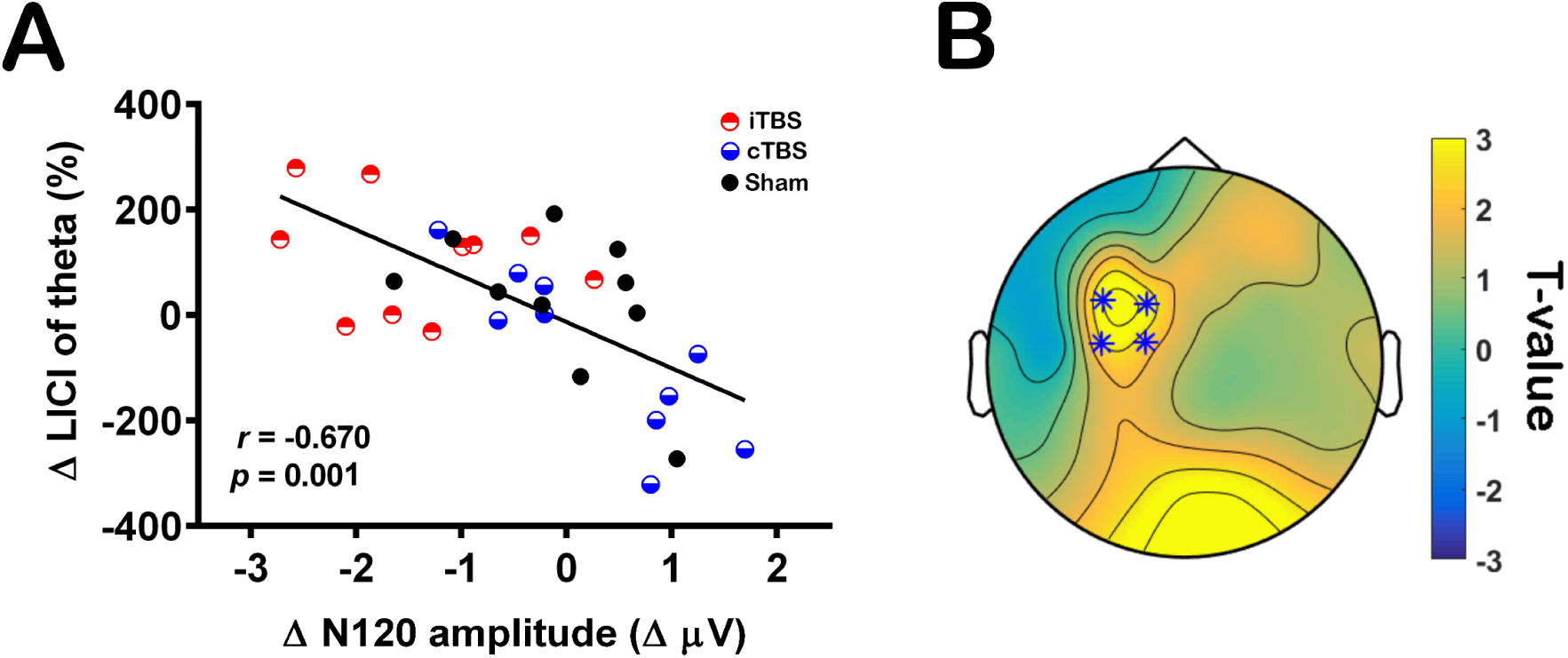
Theta-burst stimulation (TBS) induced long-interval intracortical inhibition (LICI) of transcranial magnetic stimulation (TMS)-evoked oscillations. (A) Correlation between TBS induced N120 amplitude change and change in LICI of theta frequency. (B) Intermittent TBS (iTBS) induced change in LICI of theta frequency (5 – 7 Hz; 50 – 250 ms) across the scalp. Asterisks on topoplot indicate significant clusters between pre- and post-iTBS (cluster-based statistics, \**p* < 0.01).

#### 3.2.3 Global analysis of the effect of TBS on LICI of TMS-evoked oscillations

One significant positive cluster was observed in LICI of theta (*p* = 0.007; increase LICI from pre to post-iTBS) but not in gamma band. Topographical representation revealed the difference in LICI of theta at the site of stimulation (Fig. 5B). No significant difference (all *p* > 0.05) was observed with cTBS or sham in any frequency band across the scalp.

Between stimulation conditions, one significant positive cluster in Δ LICI theta was found at the prefrontal region between iTBS and cTBS (*p* = 0.013; Δ LICI theta post-iTBS larger than post-cTBS). No other significant cluster was found between other conditions.

### 3.3 Control analysis

To ensure that the baseline correction window used for single pulse data analysis did not influence the results, we repeated the analysis using a window closer to the pulse (-500 to - 10 ms).

#### TEPs

For ROI analysis, two-way repeated measures ANOVA yielded a significant interaction (*F*_2,18_ = 5.967, *p* = 0.010) in N120 amplitude. Globally, cluster-based statistics across space showed one significant negative cluster (*p* = 0.001) at N120, and one positive cluster (*p* = 0.015) at P200 following iTBS. One significant negative cluster was observed for the change in N120 (Δ N120) [iTBS > cTBS (*p* = 0.0004); iTBS > sham (*p* = 0.032)], and one positive cluster each for the change in P200 (Δ P200) [iTBS > cTBS (p = 0.015); iTBS > sham (*p* = 0.017)].

#### TMS-evoked oscillations

For ROI analysis, two-way repeated measures ANOVA yielded a significant interaction (*F*_2,18_ = 7.857, *p* = 0.004) in TMS-evoked theta power. Globally, one significant positive cluster in the change in theta power (Δ theta) was found between iTBS and cTBS (*p* = 0.008) on the contralateral site of stimulation.

## 4. Discussion

Results of this study have demonstrated short-term plastic changes by TBS delivered to the left DLPFC. Most notably, iTBS increased N120 TEP peak amplitude, TMS-evoked theta power and LICI of TMS-evoked theta oscillations, while cTBS showed a decrease in these measures. In addition, changes in N120 amplitude following TBS correlated with change in LICI of theta oscillations, suggesting N120 may be linked to TBS-induced modulation of cortical inhibition.

In the current study, we showed that iTBS over DLPFC increased N120 amplitude, whereas we could not find evidence for a similar effect with cTBS. This is the first study to assess the effects of TBS / rTMS on DLPFC TEPs, however, studies from the motor cortex have found that TEPs with similar latencies are also modulated. For example, 1 Hz rTMS, which has been known to decrease cortical excitability, applied to the primary motor cortex increased TEPs with a latency of 100 ms (i.e. N100) in healthy individuals (Casula et al., 2014), while cTBS resulted in decreased N100 (Harrington et al., 2015, Huang et al., 2015). Recently, Harrington and colleagues (2015) have demonstrated effects of TBS on cerebellum using TMS-EEG over motor cortex, and found increased N100 amplitude following iTBS and decreased N100 amplitude following cTBS (Harrington et al., 2015), which in part, is similar to the finding of the current study. However, TBS effects were projected through cerebello-thalamo-cortical pathway in the above-mentioned study, and the parameters of the stimulation used were also different (30 Hz, 80–90*%* rMT). Even though similar results have been seen in different brain regions, it remains unclear whether N100 of M1-TEP and N120 of the DLPFC-TEP are analogous responses, and further studies are needed to characterise this negative potential. In addition to the increase in N120 amplitude, we also observed an increase in P200 amplitude following iTBS. Currently, the origin of this late TEP is still unknown, but changes have been observed following tDCS (Pellicciari et al., 2013) and PAS (Huber et al., 2008, Veniero et al., 2013).

We found an increase in TMS-evoked theta oscillations following iTBS in the prefrontal region, and to a lesser extent, a decrease with cTBS. Furthermore, iTBS also resulted in an increase in LICI of TMS-evoked theta (a paired-pulse measure of cortical inhibition (Daskalakis et al., 2008, Garcia Dominguez et al., 2014)), suggesting TBS can modulate cortical inhibition in the stimulated region. The change in LICI of theta correlated with change in N120 amplitude, suggesting that LICI of theta and N120 amplitude in DLPFC may reflect similar underlying mechanisms. Additional pharmacological studies are required to establish whether N120 amplitude and LICI of theta oscillations in DLPFC are specifically sensitive to changes in GABA_B_-mediated inhibition.

Our finding of TBS-induced changes in theta oscillations is broadly consistent with the effect of TBS on cognitive performance, especially working memory (WM), where iTBS over DLPFC increases WM performance (Hoy et al., 2015) and cTBS decreases performance (Schicktanz et al., 2015). Theta frequency is thought to reflect active operations of the cortex, such as memory encoding, retention and retrieval (Jacobs et al., 2006, Itthipuripat et al., 2013, Cavanagh et al., 2014), suggesting these changes in cognitive performance may result from changes in the capacity to generate theta oscillations.

Contrary to our hypothesis, we did not observe any change in TMS-evoked gamma oscillations. Gamma oscillations reflect coordinated neuronal activity, and are implicated in spike-timing-dependent plasticity (Nyhus et al., 2010). In an animal study, when equivalent repeats of the two TBS paradigms were applied to rat cortex, iTBS but not cTBS caused a lasting increase in gamma power of the EEG recorded from the frontal cortex (Benali et al., 2011). It should be noted that TMS-evoked oscillations may not necessarily reflect an increase in gamma power of the cortex, and we did not measure resting EEG data to confirm this finding.

No significant changes in TEPs could be detected following cTBS. This is likely to be due to individual variability to TBS, which may arise from the activation of different interneuronal networks across individuals (Hamada et al., 2013). In addition, 120% rMT was used to probe TBS-induced changes. It has been described that the magnitude and the consistency of MEP suppression induced by cTBS were greatest when probed using stimulus intensities at or above 150% rMT, while facilitation of MEPs following iTBS was strongest and most consistent at 110% rMT (Goldsworthy et al., 2016). Whether a similar relationship exists in the DLPFC requires further investigation. Vernet and colleagues (2013) have demonstrated a decrease in TMS-evoked oscillations in low frequencies (theta and alpha) in the motor cortex following cTBS (Vernet et al., 2013), which is somewhat consistent with our findings. Overall, the changes in N120 and TMS-evoked theta were significantly different between iTBS and cTBS, consistent with differential changes in MEP amplitude observed in motor cortex.

Several limitations of the present study should be acknowledged. It is possible that the TMS click sound was not completely suppressed by the application of white noise. In addition, the click of the coil transmitted via bone conduction could not be avoided in this study. Even though it is unlikely that N100-P180 components are induced by auditory effects alone (ter Braack et al., 2015), any remaining effects of auditory-evoked potentials cannot be ignored for proper inference of physiological meaning of the TBS. It is, however, worth noting that the auditory artefacts should be consistent across time, and therefore changes in TEP amplitude can be attributed to TMS-evoked neural activity. Magnetic artefacts produced by the TMS pulse can affect the measurement of evoked potentials which could confound the interpretation of EEG data. However, as we used a repeated measures, sham-controlled design it is unlikely that confounding factors would have affected the pre- and post-TBS measures unequally unless they specifically impact on the cortical plastic response to TBS. It is reassuring that we saw no significant change in cortical reactivity with sham stimulation. This study was performed on a small number of participants. A larger sample size would be required for more accurate estimation for future comparisons. Given the low number of trials collected, this may have limited the ability to detect the changes in early TEPs due to low signal-to-noise ratio. In addition, differences in LICI of TEPs were not found, which may have been impacted by a saturation of LICI at baseline preventing the assessment of increased LICI following TBS. Due to local power-line frequency at 50 Hz, analysis specific to the frequency of interest was limited, and accounted for using a wider window (30 Hz - 70 Hz), which may have resulted in the negative finding in gamma frequency. Finally, localisation of DLPFC was estimated using F3 electrode, and this could be improved by using neuronavigation.

## 5. Conclusion and future directions

This study provides preliminary evidence that TBS produces direct changes in cortical reactivity in the prefrontal cortex, in a manner similar to that seen in motor regions. We have also demonstrated that the TMS – EEG methodology can be used to study the effects of TBS in non-motor regions. This may be a useful approach to pre-clinically optimise stimulation paradigms prior to the conduct of clinical trials.

The continuing optimisation of TBS protocols is imperative in order to facilitate their translation into clinical use. For example, experiments aimed at investigating the effects of change in pulse frequency and intensity are required to define optimal stimulation parameters. Methods such as those applied in this study allow the pre-clinical investigation of optimal stimulation parameters in a manner that can potentially help develop optimal methods for clinical trial applications. Furthermore, whilst the lasting effects of theta burst stimulation have been demonstrated (Nyffeler et al., 2006, Noh et al., 2012, Goldsworthy et al., 2013), further evidence is required to demonstrate the effects of repeated TBS sessions on cortical excitability in non-motor brain regions. Studying the effect of state-dependency factors on TBS-induced cortical excitability is also important. A range of studies has demonstrated the impact of factors such as prior stimulation by TMS (Lang et al., 2004) and antiepileptic medication (Fregni et al., 2006). Examination of the state of the brain prior to TBS administration may offer further advantages in terms of optimisation. A number of studies have suggested a role for matching the stimulation frequency to the specific neural frequency of the brain prior to stimulation (Jin et al., 2006, Jin et al., 2012, Leuchter et al., 2013) and this may yield interesting results in modulating cortical plasticity using TBS.

